# Telomere 3’-overhang attrition and persistent telomeric DNA damage response in failing human hearts

**DOI:** 10.1101/2025.08.12.669993

**Authors:** Bo Ye, Liang Shang, Xun Yuan, Yuyong Xu, Ying Wen, Yongying Shi, Haodong Xu, Xuejun Wang, Kenneth B. Margulies, Xiaojing Wang, Faqian Li

## Abstract

**Background:** Telomere homeostasis is critical for normal cellular and organ function, and its dysregulation is implicated in aging and chronic diseases. Although telomere length (TL) is critical for normal telomere function, its functional status can also be altered by many other factors. The organization and protective status of telomere in failing human heart (FHH) remains incompletely understood.

**Methods:** Using left ventricular tissues from patients with idiopathic dilated cardiomyopathy (IDC), ischemic heart disease (IHD), and non-failing heart (NFH), we performed a comprehensive analysis that included measurements of overall TL and 3’ overhang length in left ventricles and cardiomyocytes (CM), assessment of telomere-binding protein associations, and transcriptomic profiling through RNA sequencing.

**Results:** Although TL varied among individuals, reduced median TL was observed only in IDC, while both IDC and IHD showed an increased frequency of very short telomeres, falling below the 5th percentile, in CMs. Strikingly, significant telomere 3’-overhang attrition was detected in both disease groups and strongly correlated with elevated H2AX phosphorylated on serine 139 (γH2AX) in heart tissues, indicating DNA damage. This was accompanied by persistent activation of ataxia telangiectasia mutated (ATM) protein-mediated DNA damage responses and the formation of telomere dysfunction-induced foci (TIFs) in CMs from FHH. Concomitantly, the association of telomeres with the single-stranded telomere-binding protein, Protection of Telomeres 1 (POT1), and the double-stranded telomere-interacting protein, telomere repeat-binding factor 2 (TERF2), was markedly reduced, accompanied by an increase in association of γH2AX and ssDNA binding protein, RPA with telomeres in IDC and IHD relative to NFH, signifying telomere de-protection.

**Conclusions:** CM telomere dysfunction, characterized by 3’ overhang attrition and de-protection, is a common feature in FHH, leading to persistent DNA damage response in telomeres. Better understanding of telomere biology throughout the progression of different heart diseases to heart failure will provide more effective prevention and treatment strategies.

**GRAPHICAL ABSTRACT:** Human telomeres safeguard chromosome ends through a higher-order telomere-loop (t-loop) structure and the shelterin complex, preventing their recognition as DNA double-strand breaks (DSBs). In the failing human heart (FHH) with idiopathic dilated cardiomyopathy (IDC) and ischemic heart disease (IHD), cardiomyocyte telomeres lose protection due to disruption of the shelterin complex, driven by oxidative DNA damage from reactive oxygen species (ROS) targeting G-rich telomeric sequences. This leads to t-loop unfolding, DDR cascade activation, 3’ overhang excision, and persistent DNA damage responses with Telomere Dysfunction-Induced Foci (TIFs) formation at the telomeric regions of cardiomyocytes in FHH. The ATM-mediated DDR, along with engagement of both classical non-homologous end joining (cNHEJ) and homologous recombination (HDR) repair pathways, is integral to FHH progression. This study underscores the significance of telomere 3’ overhang attrition in the pathogenesis of FHH with IDC and IHD. Unlike changes in overall telomere length, 3’ overhang shortening directly correlates with increased DNA damage in human myocardial tissue. Preventing telomere 3’ overhang attrition and oxidative damage offers a promising avenue for attenuating heart disease progression toward heart failure. In particular, selective ATM inhibition, as opposed to ATR inhibition, presents a potential therapeutic strategy for patients with IDC and IHD. Future efforts to pinpoint deficiencies in cNHEJ and HDR pathways in failing human hearts may reveal novel targets for heart failure prevention and treatment, providing new strategies in the management of this patient population. **CMs**, cardiomyocytes; **DDR**, DNA double-strand break repair; **FHH**, failing human heart; **ROS**, reactive oxygen species; **t-loop**, telomere-loop.

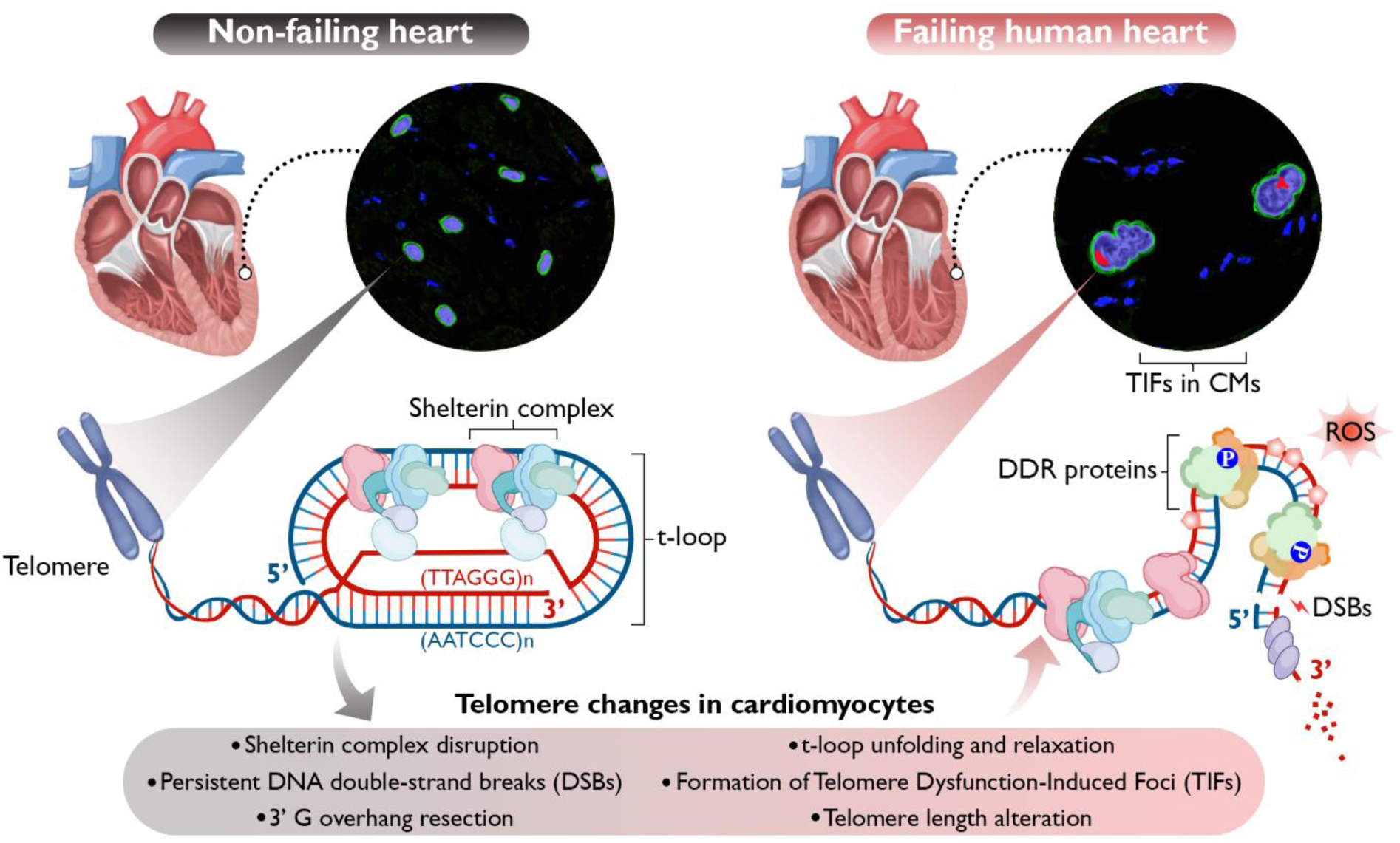

## INTRODUCTION

Mammalian chromosomes contain linear DNA with two ends that simulate double strand DNA breaks (DSBs).^1,2^ However, the telomeres located at the ends of chromosomes, characterized by linear arrays of tandemly repeated TTAGGG sequences, are effectively shielded by the shelterin complex and the telomere-loop (t-loop) structure, enabling them to evade immediate DNA damage detection and responses. This shelterin complex comprises single-stranded DNA (ssDNA) and double-stranded DNA (dsDNA) binding proteins, along with regulatory molecules, forming a terminal DNA and protein complex.^3–5^ In addition to the complimentary telomere DNA duplex, there exists a single-stranded G rich TTAGGG overhang at the 3’ end of telomeres. This overhang plays a crucial role in forming a protective higher-order t-loop structure, where the 3’ overhang strand folds back and invades a proximal region of the same chromosome end. Any attrition of the telomere 3’ overhang or shortening of the duplex can disrupt t-loop formation, leading to telomere de-protection and dysfunction.^1,6,7^ In humans, telomere duplex binding proteins, including telomere repeat-binding factor 1 (TERF1) and 2 (TERF2), along with the single-stranded interacting protein, protection of telomeres 1 (POT1), which assembles as a heterodimer with TPP1 (ACD), are essential for normal telomere protection and homeostasis.^3–5^

Telomere length (TL) and composition of telomere-associated proteins are tightly regulated and undergo dynamic changes during the cell cycle, development, and aging, as well as in response to a variety of physiologic challenges and pathologic stresses.^8^ During each cell cycle, telomeres of dividing cells naturally shorten due to the end replication problem, ultimately leading to replicative senescence.^9^ This telomere attrition in cells has been observed to accelerate during periods of stress and in a variety of human diseases.^8,10^ Interestingly, terminally differentiated cells such as cardiomyocytes (CMs), neurons, and skeletal muscles, which have limited proliferative activity, also experience telomeric DNA shortening during aging or under stresses. The mechanisms by which non-dividing cells lose their telomere sequences are not well understood. Telomere shortening can cause telomere de-protection, but de-protected telomeres are not necessarily short.^11^ Therefore, it is not just TL, but also telomere status that plays a crucial role in modulating cellular function, metabolism, and organismal aging. The telomere status in chronic diseases, especially in failing human hearts (FHHs), has not been clearly defined. Here, we aim to determine TL and assess its protective status in FHH with idiopathic dilated cardiomyopathy (IDC) and ischemic heart disease (IHD) in comparison to those in non-failing human hearts (NFHs).

## METHODS

Detailed methods are provided in the Supplemental Materials.

### Human Hearts

Heart tissues from the left ventricular free wall of 63 human hearts, including 22 IDCs, 21 IHDs and 20 NFHs from patients without a history of cardiac diseases, were procured using a standardized protocol.^12–14^ Clinical data pertaining to the patients are summarized in Table 1 and listed in Supplemental Table S1. All experiments involving human tissues were performed under protocols approved by Institutional Review Boards at the University of Pennsylvania and the Gift-of-Life Donor Program (Pennsylvania, USA), University of Minnesota, and University of Texas Health Center at San Antonio.

**Table 1.**
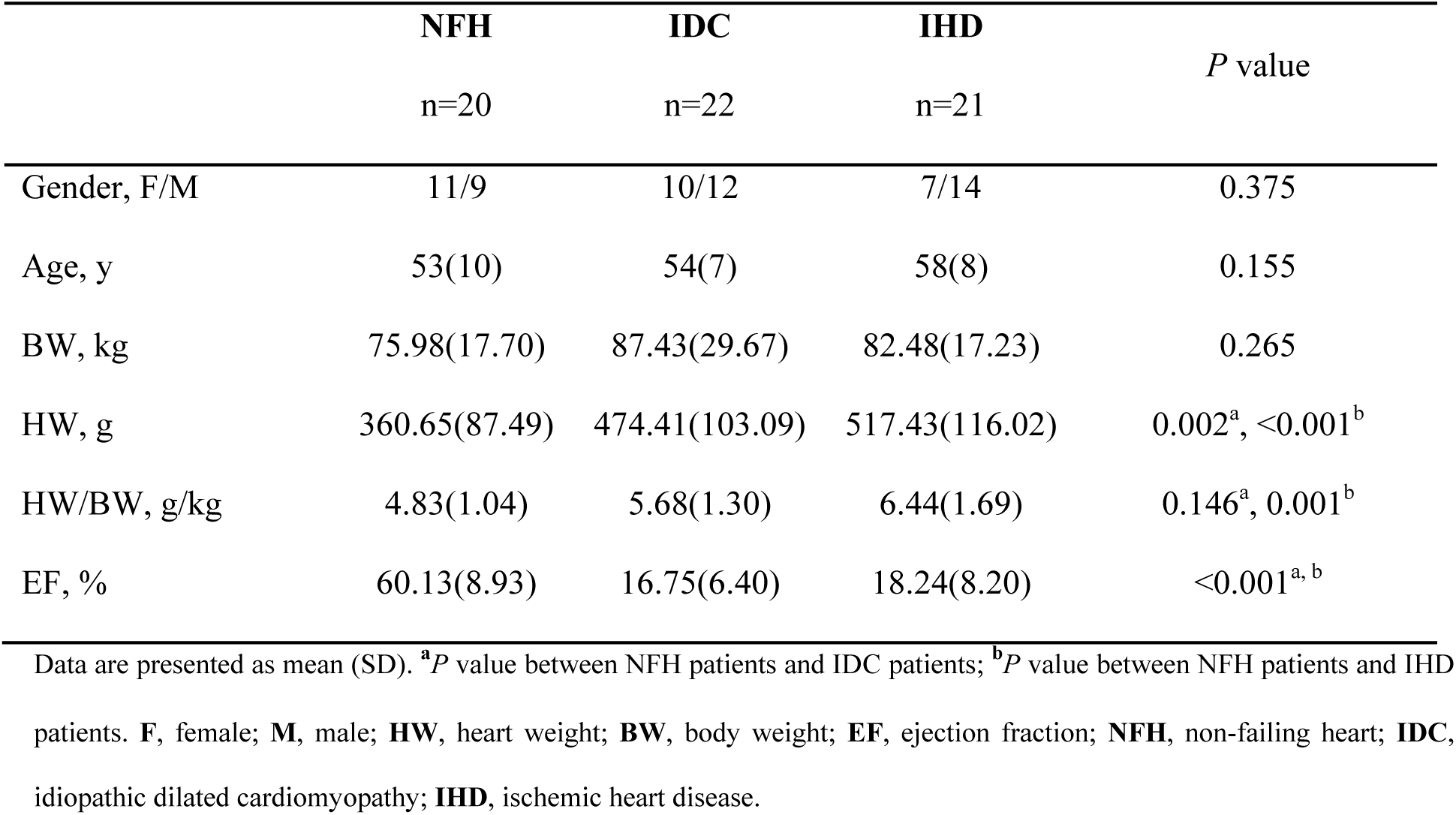
Patient baseline characteristics of heart tissues used in this study.

|  | **NFH**<br>n=20 | **IDC**<br>n=22 | **IHD**<br>n=21 | *P* value |
| --- | --- | --- | --- | --- |
| Gender, F/M | 11/9 | 10/12 | 7/14 | 0.375 |
| Age, y | 53(10) | 54(7) | 58(8) | 0.155 |
| BW, kg | 75.98(17.70) | 87.43(29.67) | 82.48(17.23) | 0.265 |
| HW, g | 360.65(87.49) | 474.41(103.09) | 517.43(116.02) | 0.002<sup>a</sup>, <0.001<sup>b</sup> |
| HW/BW, g/kg | 4.83(1.04) | 5.68(1.30) | 6.44(1.69) | 0.146<sup>a</sup>, 0.001<sup>b</sup> |
| EF, % | 60.13(8.93) | 16.75(6.40) | 18.24(8.20) | <0.001<sup>a, b</sup> |

Data are presented as mean (SD). <sup>a</sup>*P* value between NFH patients and IDC patients; <sup>b</sup>*P* value between NFH patients and IHD patients. **F**, female; **M**, male; **HW**, heart weight; **BW**, body weight; **EF**, ejection fraction; **NFH**, non-failing heart; **IDC**, idiopathic dilated cardiomyopathy; **IHD**, ischemic heart disease.

### Statistical Analysis

Categorical variables were expressed as counts and percentages and were compared across groups using the Chi-squared test followed by post hoc cellwise tests using adjusted standardized residuals when the expected count for each cell was five or more, otherwise Fisher’s exact test was used. Continuous data were reported as mean and standard deviations (SD) or median. Statistical analyses were conducted using SPSS19.0 or GraphPad 9.0 software. Standard unpaired Student’s t-tests, Mann-Whitney *U* test or Median test (two-tailed; *P*<0.05) were used for comparing two groups when appropriate. For multiple-comparison analysis, One-way ANOVA with Bonferroni’s or Dunnett’s post-test correction was applied for single-factor analysis, and two-way ANOVA followed by post hoc comparisons was used for two-factor analysis when appropriate (*P*<0.05). Kruskal-Wallis test was applied to compare medians among more than two groups, followed by Mann-Whitney post hoc tests for pairwise comparisons, with a Bonferroni correction for multiple hypothesis testing. The correlation between two variables was assessed using the SPSS curve estimation function. The goodness-of-fit of a curve was graphically and numerically evaluated based on residuals plotting, prediction bounds and goodness-of-fit statistics. Additionally, the coefficient of determination, R-square (R^2^), is utilized to describe the strength of the linear and non-linear relationship between the two variables.

### Data availability

All data supporting the findings of this study are available within the article and its supplemental information files, including the supplemental Supporting Data Values file. Additional data, analytic methods, and study materials related to this paper will be made available to other researchers for purposes of reproducing these results or replicating these procedures upon reasonable request to the corresponding author.

## RESULTS

### TL varies in human hearts and very short telomeres are present in IDC and IHD

TL in human hearts varied among individuals, as determined by telomere restriction fragment (TRF) analysis using genomic DNA (Figure 1A). The average TRF lengths for NFH, IDC and IHD were 8862, 8555 and 8522 base pairs (bp), respectively. No statistically significant differences were detected among the three groups by TRF analysis (Figure 1B). Similarly, there was no difference observed among the three groups when examining the relative average TL based on telomere signals normalized to a single-copy gene 36B4 (T/S ratio) by quantitative PCR (qPCR) (Figure 1C). The median TL of CMs was shorter in IDC (*P*<0.001), with no apparent changes observed in IHD (*P*=0.130), as demonstrated by Q-FISH (Figure 1D). Contrarily, the mean TL of CMs was longer in IHD, with no changes found in IDC (NFH, 1.36×10^6^; IDC, 1.34×10^6^, *P*=0.988; IHD, 1.87×10^6^, *P*<0.001). Additionally, there was an increase in the percentage of very short telomeres, falling below the 5th percentile of telomere integrated intensity, in both IDC (15.15%, *P*<0.001) and IHD (7.67%, *P*=0.005) compared to NFH (5.02%). Furthermore, IHD contained more telomeres above the 95th percentile of normal telomere distribution compared to NFH (*P*<0.001), exhibiting a non-normal pattern with a higher concentration of data points in both the 5th and 95th percentiles. These results indicate that although the overall average or median TL determined by established methods may not change, there are significantly more outliers in the TL of CMs from FHH compared to those from NFH.

**Figure 1.**
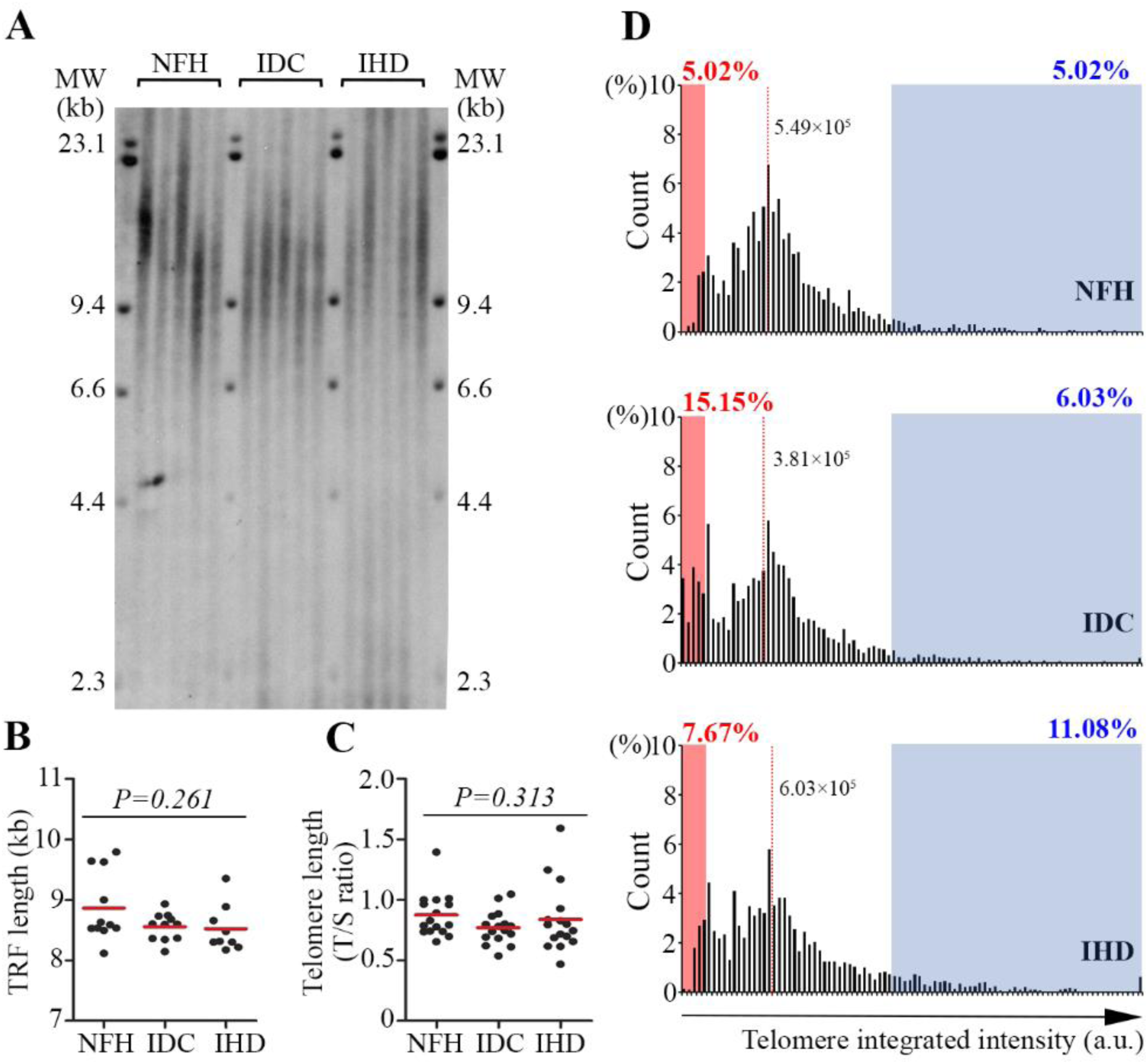
Very short telomeres increase while total telomere length varies in failing human hearts. **A**, Representative images from telomere restriction fragment (TRF) analysis via Southern Blot illustrate telomere length (TL) in non-failing heart (NFH), idiopathic cardiomyopathy (IDC) and ischemic heart disease (IHD). **B**, TRF quantification reveals no significant differences in average TL among NFH, IDC, and IHD. NFH: n=11; IDC: n=11; IHD: n=9. **C**, Average TL, assessed by telomere signals normalized to the single-copy gene 36B4 (T/S ratios) via qPCR, reveals no differences among the three groups. A total of 16 samples for each group (NFH, IDC, and IHD) were analyzed. **D**, Histograms of telomere integrated intensity by quantitative fluorescence hybridization (Q-FISH) with Alexa 488-labeled telomere peptide nucleic acid (PNA) probes depict similar median TL in cardiomyocytes (CMs) from NFH and IHD, but a shorter TL in IDC. Red dotted lines indicate the median TL in CMs for each group. Red rectangles highlight a higher percentage of very short telomeres below the 5th percentile in IDC and IHC. Blue rectangles denote a higher percentage of telomere integrated intensity above the 95th percentile in IHD. 75-100 CMs were randomly selected for each group from 11 NFH, 12 IDC, and 10 IHD samples.

### Truncation of 3’ overhang is a common feature of IDC and IHD

We assessed the length of the 3’ overhang through dot blots using native genomic DNA and biotinylated (CCCTAA)_3_ telomere probes. The 3’ overhang signals were significantly weaker in IDC and IHD compared to NFH. In contrast, there was less variation among the three groups when analyzing the denatured DNA for total TL (Figure 2A and B). To confirm that the telomere 3’ overhang signal under native conditions reflects the length of 3’ overhang, we selected samples with strong 3’ overhang signals from IDC, IHD, and NFH groups for exonuclease digestion. Exonuclease I (EXO I) removes 3’ ssDNA, while exonuclease VII (EXO VII) removes both 5’ and 3’ ssDNA, thereby eliminating 3’ overhang signals. Exonuclease III (EXO III), a dsDNA-specific 3’ exonuclease, does not affect telomere 3’ overhang signals. This method is well established for detecting and verifying 3’ overhang signals and has been used in previous studies.^15,16^ After digestion with single-stranded 3’ exonuclease EXO I or single-stranded 5’ and 3’ exonuclease EXO VII, the telomere 3’ overhangs were abolished. Importantly, dsDNA 3’ specific exonuclease EXO III did not affect the detection of telomere 3’ overhangs (Figure 2C). Our results confirm that 3’ overhangs can be successfully detected through dot blotting under non-denaturing conditions, and that the attrition of these overhangs is a prevalent feature in FHH. Our data demonstrate, for the first time, that 3’ overhang truncation is involved in heart failure.

**Figure 2.**
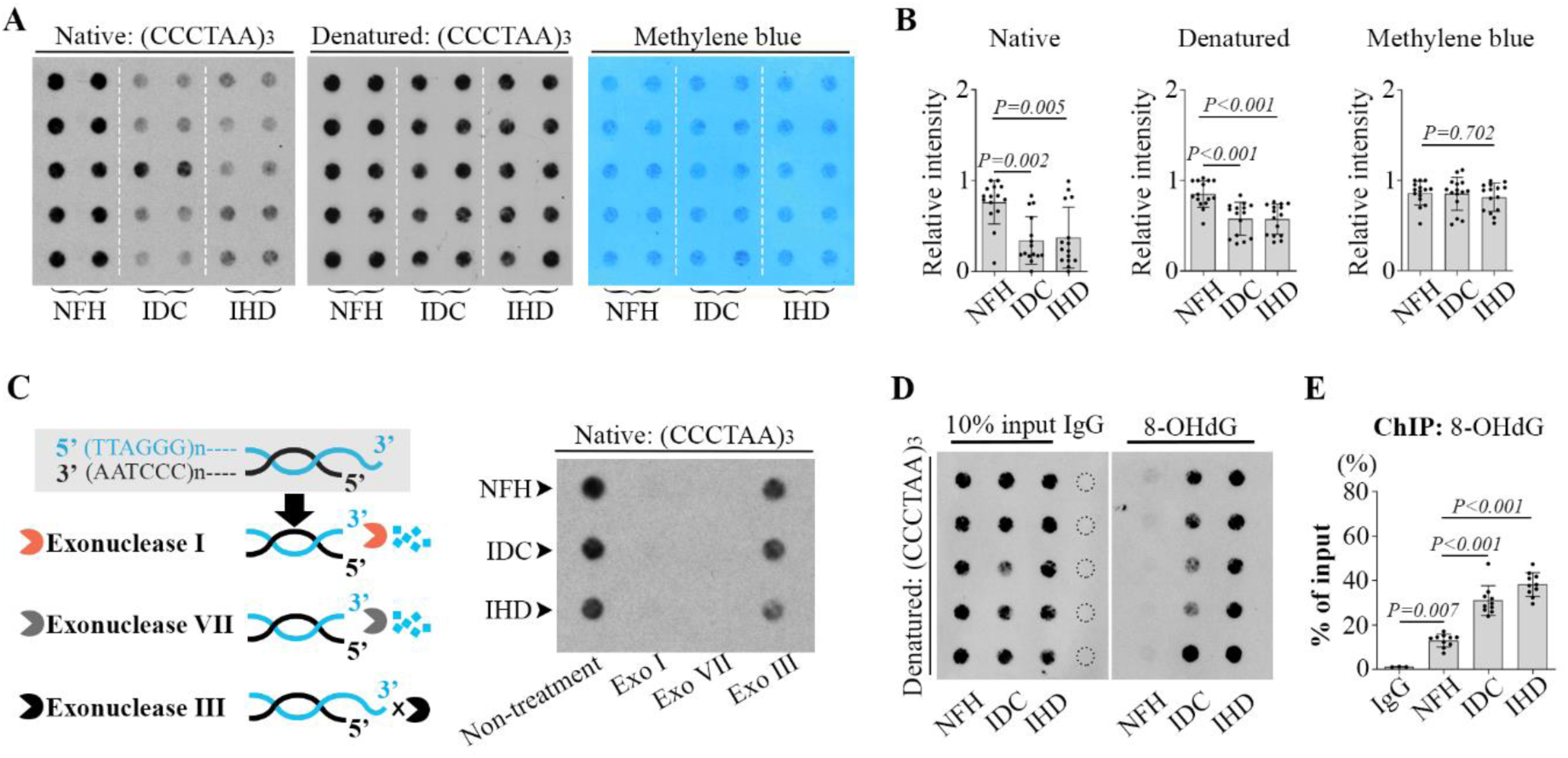
Truncation of 3’ overhang is a common feature in failing human hearts. **A**, Dot-blots of native and denatured genomic DNA hybridized to biotinylated telomere (CCCTAA)_3_ oligonucleotide probes reveal much weaker signals under native conditions in idiopathic cardiomyopathy (IDC) and ischemic heart disease (IHD) than in non-failing heart (NFH). Methylene blue staining serves as a loading control. **B**, Dot-blot signal intensities were quantified using ImageJ and normalized to NFH samples. NFH, n=15; IDC, n=15; and IHD, n=15. **C**, Exonuclease treatment for 3’ overhang detection. Schematic illustrations (left) show exonuclease directionality on the 3’ DNA end. Dot blot (right) shows that exonuclease I and VII remove 3’ single-stranded DNA (ssDNA), thereby eliminating 3’ overhang signals. Exonuclease III, a double-stranded DNA (dsDNA) specific 3’ exonuclease, has no impact on telomere 3’ overhang signals. **D** and **E**, Chromatin immunoprecipitation (ChIP) followed by dot blots (D) and qPCR (E) reveals 8-OHdG enrichment at telomere regions in IDC and IHD. ChIP-dot blot analysis included 5 samples each from NFH, IDC, and IHD, while ChIP-qPCR involved 10 hearts from each group (NFH, IDC, and IHD).

Telomeres are particularly susceptible to oxidative DNA damage caused by reactive oxygen species (ROS) due to the presence of their G-rich sequence leading to the formation of 8-hydroxydeoxyguanosine (8-OHdG).^17,18^ We hypothesized that oxidative stress could be contributing to telomere 3’ overhang shortening in FHH. We performed ChIP using an antibody against 8-OHdG, followed by dot blotting of denatured pull-down chromatin with biotinylated (CCCTAA)_3_ telomere probes and qPCR with Tel F and Tel R primers. The results showed a robust enrichment of 8-OHdG in telomeres from IDC and IHD compared to NFH (Figure 2D and E).

### DNA damage increases in CMs, but not in non-CMs of FHH

3’ G-rich overhang truncation can disrupt t-loop formation, leading to telomeres being recognized as DSBs and triggering DNA damage responses.^2^ Histologically, CMs in FHH with IHD and IDC show enlarged nuclei with irregular contours and high DNA content (Figure 3A and Supplemental Figure S2), which are signs of cells experiencing DNA damage.^19^ In FHH, myocardial DNA damage appears to be an independent predictor of heart failure outcomes, regardless of the duration of heart failure.^20^ Using the DSB marker γH2AX, we found higher percentages of γH2AX-positive CMs and more γH2AX foci per CM in IHD compared to IDC and NFH. While the percentage of γH2AX-positive CMs was similar between IDC and NFH, IDC had significantly more γH2AX foci per CM compared to NFH (Figure 3A-C). Notably, γH2AX foci were rarely detected in non-CMs lacking the CM-specific nuclear marker PCM-1(Figure 3A). We also measured the size of γH2AX foci in γH2AX-positive CMs from IDC, IHD, and NFH, revealing high variability with larger average volumes of γH2AX foci in IHD compared to IDC and NFH (Figure 3D). Western blots further confirmed increased γH2AX levels, despite variations observed among individual samples, and elevated γH2AX/H2AX ratios in FHH relative to NFH. Additionally, we observed activation of ATM but not ATR in FHH, as indicated by the pATM/ATM ratio, which was significantly elevated in IHD relative to NFH (Figure 3E and Supplemental Figure S4A). To assess the impact of potential explanatory variables on γH2AX levels, curve estimation regression was performed to investigate the correlation between telomere 3’ G-rich overhang lengths, as determined through native telomeric DNA dot blots and presented as integrated density, and relative γH2AX protein levels in human heart tissues (Table S4). One of the best-fitting curves showed a robust negative correlation in a quadratic model between telomere 3’ G-rich overhang lengths and relative γH2AX protein levels in the same heart samples from individual patients (Figure 3F). This suggests that human hearts with shorter 3’ G-rich overhangs exhibit higher γH2AX levels. Additionally, regression analyses between telomere overhang lengths and clinical characteristics showed that patients with shorter 3’ G-rich overhangs tend to have a higher body mass index (BMI). However, there was no significant impact observed on left ventricular ejection fraction (LVEF), creatinine levels, or heart weight to body weight ratios (H/B ratios) (Figure S3 and Table S4).

**Figure 3.**
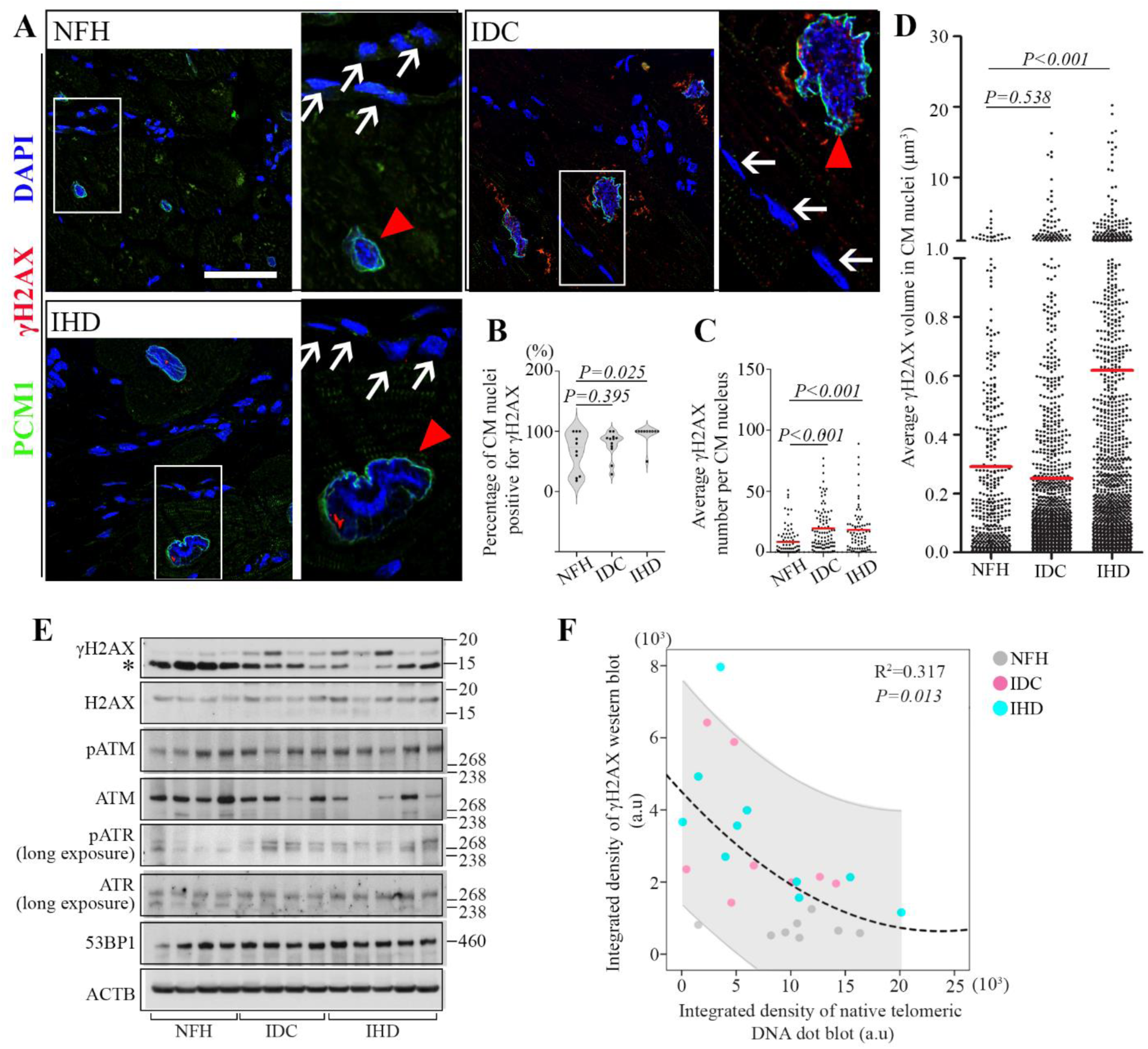
Elevated DNA damage in cardiomyocytes of failing hearts. **A**, Representative confocal images show increased levels of the DNA double-strand break marker, γH2AX (red), within the nuclei of cardiomyocytes (CMs, red triangles) positive for the CM-specific nuclear marker, PCM1 (green), in idiopathic cardiomyopathy (IDC) and ischemic heart disease (IHD) compared to non-failing heart (NFH). Non-CMs, negative for PCM1, are indicated by white arrows. Magnified views of the highlighted regions (white rectangles) are provided to the right. Lipofuscin granules appear as red signals outside the nucleus in perinuclear regions. Scale bar, 50μm. **B** and **C**, The percentage of γH2AX-positive CMs (B) is elevated in IHD, and the average number of γH2AX foci per CM (C) is higher in IDC and IHD compared to NFH. 75-98 CMs were randomly selected for each group from 10 NFH, 12 IDC, and 10 IHD samples. **D**, γH2AX foci exhibit size variation, with the average volume of γH2AX foci (red bars) being larger in γH2AX-positive CMs from IHD than those from IDC and NFH. 50-80 γH2AX-positive CMs were randomly selected for each group from 11 NFH, 12 IDC, and 10 IHD samples. **E**, Western blot analysis reveals increased levels of γH2AX and pATM, but not pATR, in failing hearts. The ratios of γH2AX/H2AX and pATM/ATM, but not pATR/ATR, measured by ImageJ are significantly elevated in IDC and IHD relative to NFH (see Supplemental Figure S3A). p53 binding protein 1 (53BP1) and beta actin (ACTB) serve as loading controls. * indicates nonspecific bands around 15 kDa (γH2AX/H2AX is around 17 kDa). A total of 8 NFH, 8 IDC, and 10 IHD samples were analyzed. **F**, A strong negative correlation is observed between γH2AX protein levels and telomeric 3’ G-rich overhang lengths in heart tissues, as indicated by the quadratic model. R² represents the coefficient of multiple determination. The grey shaded area represents the 95% confidence interval.

### Larger γH2AX foci colocalize with pATM but not ATR and are identified in CMs from IDC and IHD

We observed two types of γH2AX foci in human CM nuclei. One type, small to medium-sized, is evenly distributed throughout the CM nuclei in both NFH and FHH. The other type tends to aggregate, cluster, or merge into larger foci, often exceeding 1 μm^3^ under our observation and measurement settings (Figures 4, 5 and 6, videos S2-S4), and is predominantly present in FHH. Using confocal microscopy with 3D reconstruction, we found that pATM is more likely to colocalize with larger, aggregated γH2AX foci rather than the smaller, scattered ones (Figure 4A-B). Additionally, p53 binding protein 1 (53BP1), a downstream substrate of ATM kinase, was observed to colocalize with larger clustered damage foci, while pATR did not show colocalization with either the smaller or larger γH2AX foci (Supplemental Figure S4B). The percentage of CMs with pATM-positive γH2AX foci was significantly higher in IDC and IHD than in NFH (NFH, 2.02%; IDC, 23.52%; IHD, 30.87%), indicating a persistent DNA damage response and ATM activation in CMs of FHH (Figure 4C). The average size of pATM-positive γH2AX foci was 2.79 μm^3^, with 78% of them exceeding 1 μm^3^. In contrast, the average size for pATM-negative γH2AX foci was 0.14 μm^3^ (Figure 4D, left). There was no significant difference in the size of foci, whether positive or negative, for pATM within γH2AX-positive CMs among NFH, IDC, and IHD (Figure 4C, right).

**Figure 4.**
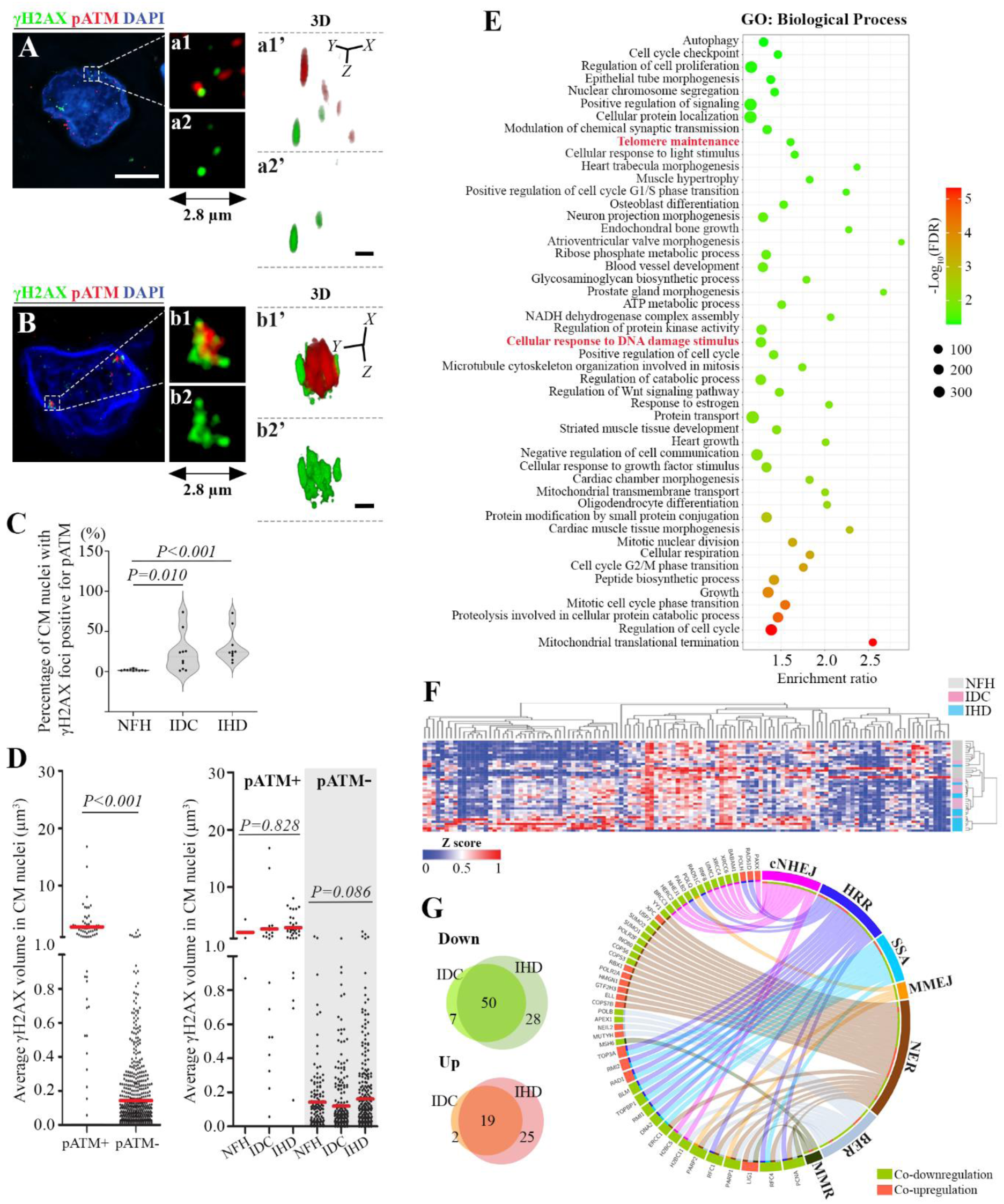
Larger γH2AX foci colocalize with pATM and are mainly identified in failing human hearts. **A** and **B**, 3D reconstructions of representative confocal images depict individual cardiomyocyte (CM) nuclei featuring scattered, smaller γH2AX foci (A) and merged, larger γH2AX foci (B), with a predominant colocalization of pATM (red) with the merged, larger γH2AX (green) foci. Zoomed-in views of the 2.8×2.8 µm² white box regions (a1 and b1) allow visual analysis and comparison of the correlation between pATM and γH2AX within distinct types of γH2AX foci (a2 and b2). These regions were reconstructed in 3D (a1’, a2’, b1’ and b2’) from 61-layer Z-stacks with 0.1-μm depth intervals for a1, a2, b1, and b2, individually. White scale bar, 10μm. Black scale bar, 1 μm. **C**, The bar graph illustrates that the percentage of CM nuclei with γH2AX foci that are also positive for pATM is significantly higher in idiopathic cardiomyopathy (IDC) and ischemic heart disease (IHD) compared to non-failing heart (NFH). Data were collected from at least 15 high-power (40×) fields for each heart. A total of 10 samples for each group (NFH, IDC, and IHD) were analyzed. **D**, The left graph shows that the average volume of γH2AX foci positive for pATM (pATM+) is notably larger than that of γH2AX foci negative for pATM (pATM-). However, there is no difference in γH2AX size among NFH, IDC, and IHD wthin pATM-positive or negative foci (right graph). Red bars represent mean of each group. 5-50 CMs positive for both γH2AX and pATM were randomly selected for each group from 10 NFH, 10 IDC, and 10 IHD samples. **E**, Gene Ontology (GO) analysis reveals that telomere maintenance and cellular response to DNA damage stimulus are among the top 45 most enriched GO terms in IDC and IHD. The size of the dots indicates the number of differentially expressed genes (DEGs) enriched in a GO term. **F**, A heatmap of hierarchically clustered, normalized, and z-scaled expression values of 131 DEGs of IDC and IHD involved in the Reactome DNA repair and damage response pathway (R-HSA-73894). **G**, Venn diagrams show the number of overlapping downregulated or upregulated DEGs, as well as the uniquely expressed DGEs, that are involved in the DNA repair and damage response pathway in IDC and IHD. The circle plot illustrates co-downregulated (green) and co-upregulated (red) DEGs in the DNA damage response for IDC and IHD, enriched in different DNA damage repair mechanisms. cNHEJ: classical nonhomologous end joining; HRR: homologous recombination repair; SSA: single strand annealing; MMEJ: microhomology-mediated end joining; NER: nucleotide excision repair; BER: base excision repair; MMR: mismatch repair.

**Figure 5.**
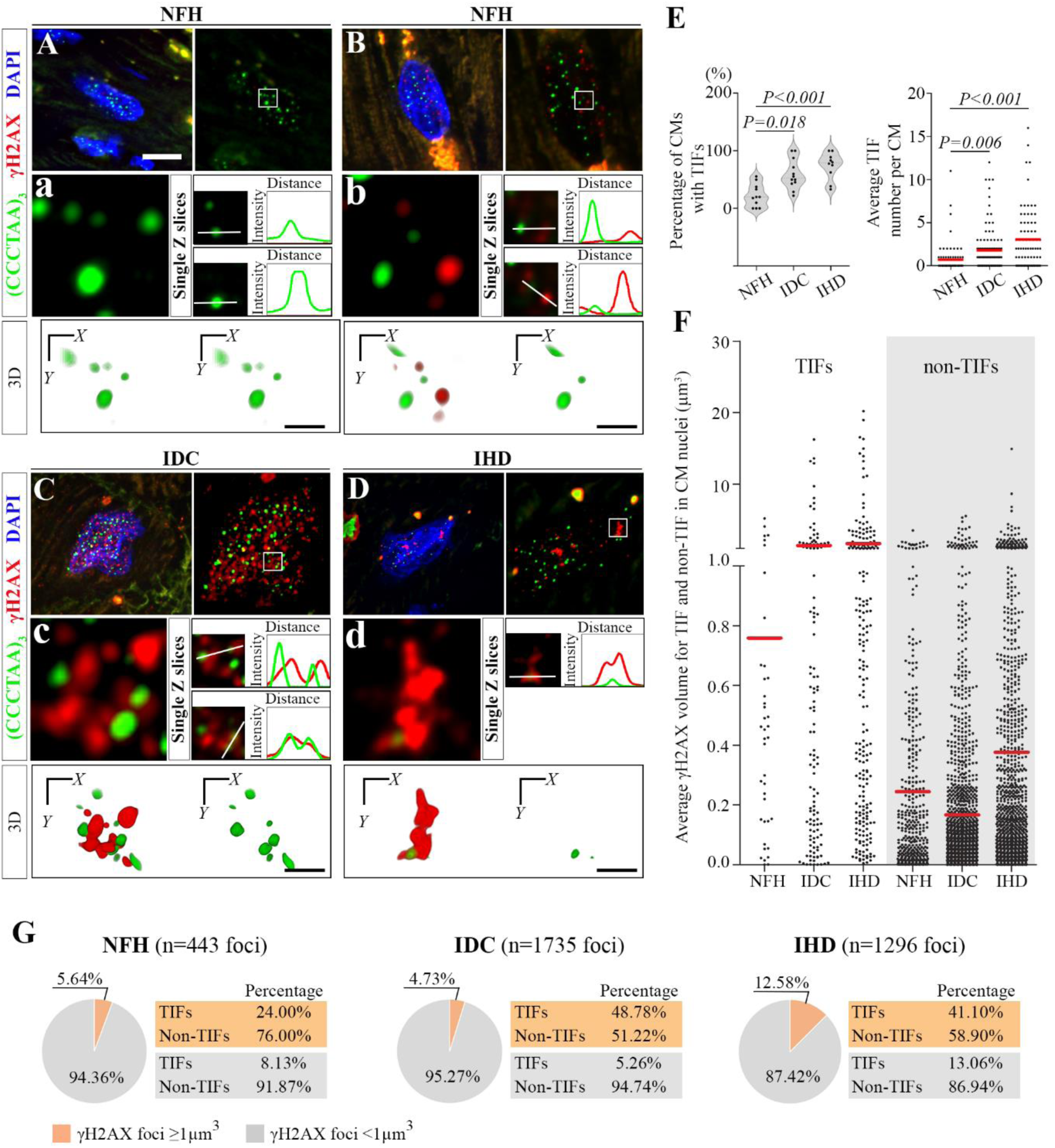
Larger γH2AX foci preferentially colocalizing with telomeres represent Telomere Dysfunction-Induced Foci (TIFs). **A-D**, Immunofluorescent (IF) and fluorescent in-situ hybridization (FISH) analysis to detect TIFs in cardiomyocytes (CMs). γH2AX (red) was detected by IF, and telomeres by FISH using a telomere probe ([CCCTAA]_3_, green). Representative maximum-intensity-projection images of randomly selected CMs from each group were obtained through confocal microscopy and deconvolution using Huygens Professional software. Zoomed-in areas (white squares, 2.8×2.8 µm) with γH2AX and telomere signals are shown at higher magnification in panels a-d. Graphs quantify γH2AX and telomere signals in regions of interest (white lines), illustrating signal colocalization and fluorescence intensity profiles using single-slice confocal images from Z-stacks of the zoomed-in areas. 3D reconstruction images were generated for the zoomed-in areas (a, b, c, and d) using 61-layer Z-stacks with 0.1-μm depth intervals. The online videos present a 360-degree rotational 3D view of these zoom-in areas (Supplemental Videos S1-4). CM nuclei in non-failing heart (NFH) are negative for γH2AX (A) or contain small γH2AX foci (B) that do not colocalize with telomeres. In idiopathic cardiomyopathy (IDC) and ischemic heart disease (IHD), larger γH2AX foci colocalize with telomeres, forming a cluster with varying strengths of telomere signals in IDC (C, Supplemental video S3) and colocalize with telomeres with an average telomere signal strength in IHD (D, Supplemental video S4), respectively. White bar, 10 µm; black bar, 1 µm. **E**, The percentage of CM nuclei with TIFs and the average TIF numbers per CM nucleus are elevated in IDC and IHD compared to NFH. For A-E, 75-100 CMs were randomly selected for each group from 11 NFH, 12 IDC, and 10 IHD samples. **F**, The average volume of γH2AX foci for TIFs is larger than that for non-TIFs (*P*<0.001). TIFs and non-TIFs were automatically determined, and their γH2AX foci volume was measured by Huygens Professional using 3D images. Red bars represent the average volume of γH2AX foci for TIFs and non-TIFs of each group. **G**, The percentage of TIFs in small (<1µm^3^) and big (≥1µm^3^) γH2AX foci was calculated in γH2AX-positive CMs and compared among CM nuclei from NFH, IDC and IHD. For analysis in F and G, 50-80 γH2AX-positive CMs were randomly selected for each group from 11 NFH, 12 IDC, and 10 IHD samples.

**Figure 6.**
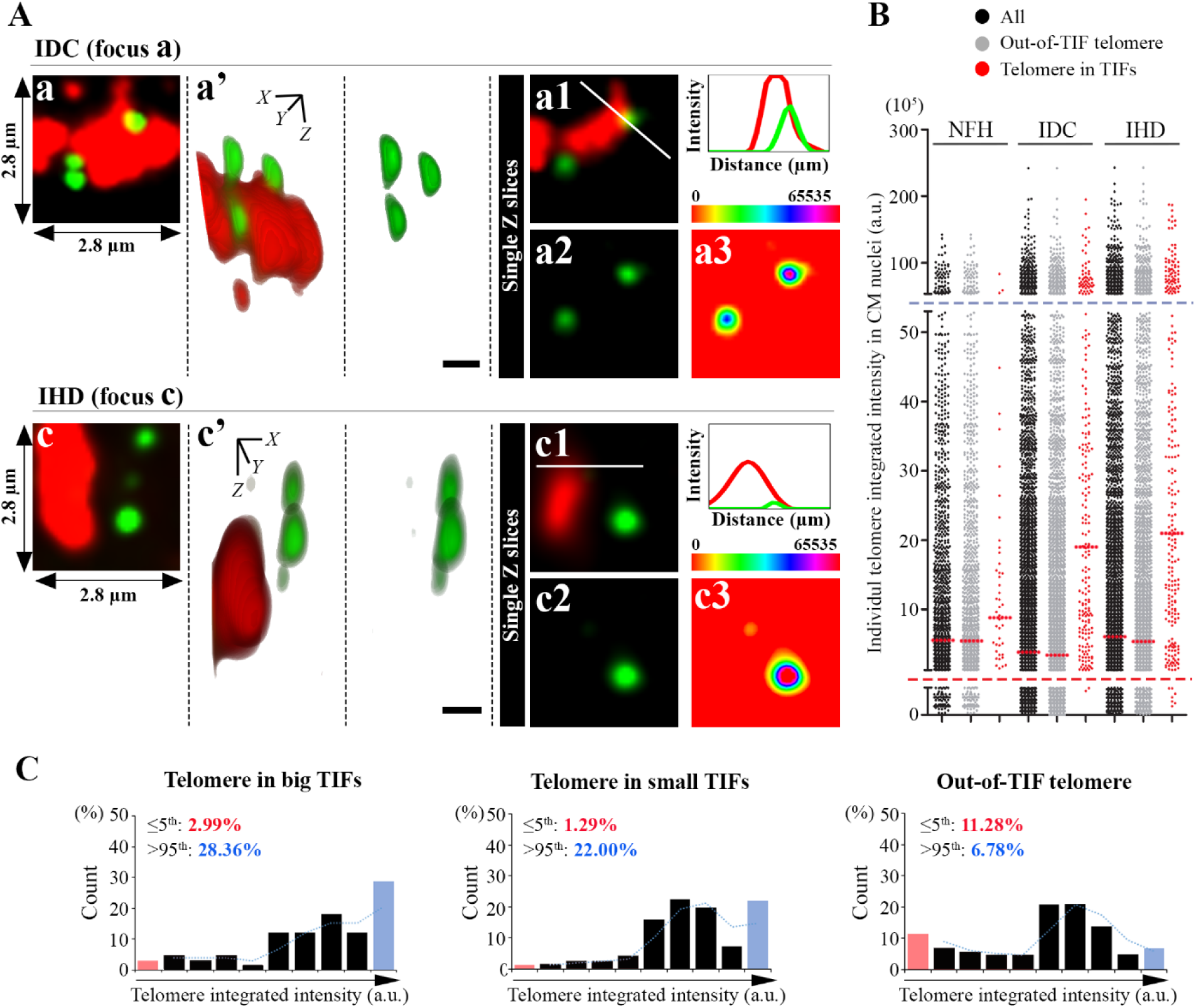
Telomeres in Telomere Dysfunction-Induced Foci (TIFs) are not necessarily shorter. **A**, Detailed 3D analysis of foci a and c from maximum-intensity-projection images (Supplemental Figure S4A) of immunofluorescence (IF) and fluorescence in-situ hybridization (FISH) for γH2AX (red) and telomeres ([CCCTAA]_3_, green) in cardiomyocytes (CMs) from idiopathic dilated cardiomyopathy (IDC) and ischemic heart disease (IHD) with large γH2AX foci (≥1µm^3^) that colocalize with telomeres. Focus a contains strong telomere signals (>95th percentile) with clustering, and c colocalizes with an extremely weak telomere signal (<5th percentile). The 3D reconstruction (a’ and c’) was generated from 61-layer Z-stacks with 0.1-μm depth intervals, covering 2.8×2.8 µm^2^ areas from foci a and c. RGB profile for signal colocalization and fluorescence intensity profiles was plotted along with the white lines in single-slice confocal images with merged (a1 and c1) γH2AX and telomere signals. Single channel for telomere signals is shown in a2 and c2. Special LUT (a3 and c3) was applied to the green channel to highlight the telomere signals. White scale bar, 10μm, and black scale bar, 1μm. **B**, Integrated intensity of individual telomeres in TIFs and those that do not colocalize with γH2AX (out-of-TIF telomeres) were measured to represent telomere length (TL) among CMs from IDC, IHD and non-failing heart (NFH). Median TL (red dotted lines) for telomeres in TIFs is longer than out-of-TIFs telomeres (Median test, *P*<0.001). Mann–Whitney test shows significant difference in telomere intensities distribution between TIFs and out-of-TIFs for CMs (*P*<0.001). The red and blue dash lines indicate 5th and 95th percentiles of telomere integrated intensity, respectively. **C**, Histogram of telomere integrated intensity by quantitative FISH (Q-FISH) with Alexa 488-labeled telomere PNA probes reveals TL variation in big TIFs, small TIFs and out-of-TIFs. 75-100 CMs were randomly selected for each group from 11 NFH, 12 IDC, and 10 IHD samples.

We re-analyzed a published RNA-seq dataset (GSE116250)^21^ of human left ventricular samples from patients with IDC and IHD, as well as NFH controls. Differentially expressed genes (DEGs) identified in IDC and IHD are listed in Tables S5 and S6. The co-regulated DEGs in IDC and IHD were further subjected to Gene Ontology (GO) analysis. Notably, the mitochondrial-related gene set showed the most significant alterations in FHH, and the cellular response to DNA damage stimulus was among the top 25 overrepresented GO terms in Biological Process ranked by the adjusted *P* value (Figure 4E). Hierarchically clustering of normalized and z-scaled expression values of 131 DEGs of FHH involved in the Reactome DNA repair and damage response pathway (R-HSA-73894) effectively segregated NFH, IDC and IHD (Figure 4F). Among the 131 DEGs, 69 genes exhibited a similar trend of changes, with 50 co-downregulated and 19 co-upregulated genes observed in both IDC and IHD (Figure 4G and Table S7). Moreover, the co-regulated DGEs integrally involved in key DNA repair mechanisms were categorized into seven pathways using Circos analysis. These pathways encompass classical nonhomologous end joining (cNHEJ), homologous recombination repair (HRR), single-strand annealing (SSA), microhomology-mediated end joining (MMEJ, also known as alt-NHEJ), nucleotide excision repair (NER), base excision repair (BER), and mismatch repair (MMR). Among these pathways, 42% of the co-regulated DGEs were exclusively dedicated to DNA double-strand break repair (DDR), executed through cNHEJ and homology-directed repair (HDR), which includes HRR, SSA, and MMEJ. Another 42% were solely implicated in NER, BER, and MMR which are major DNA repair pathways responsible for ROS-induced DNA damages. Additionally, 16% of the identified DGEs were shared between DDR-based and ROS-induced DNA damage repair categories (Figure 4G).

### Larger γH2AX foci colocalize with telomeres representing Telomere Dysfunction-Induced Foci (TIFs) that increase in IHD and IDC

While most γH2AX foci were not associated with telomeres (Figure 5A-D), there was a significant increase in the percentage of CM nuclei with telomeres positive for γH2AX, indicating TIFs, in IDC (59.48%) and IHD (73.93%) compared to NFH (26.20%). The average number of TIFs per CM also increased in FHH (Figure 5E). Comprehensive analyses using confocal microscopy, immunofluorescent (IF) labeling of γH2AX and telomere Q-FISH, followed by 3D deconvolution and reconstruction, revealed a preferential colocalization between clustered, larger γH2AX foci and telomeres (Figure 5C-D). The size differences of γH2AX foci can be challenging to determine in the Z-stack-projection images but become more apparent after 3D reconstruction (videos S1-S4). Notably, the average size of γH2AX foci for TIFs in NFH, IDC and IHD was 0.76, 1.41 and 1.70 μm^3^, respectively, and TIFs were significantly larger than non-TIFs (*P*<0.001, Figure 5F). Two-way ANOVA revealed a significant interaction between foci type (TIFs or non-TIFs) and tissue type (IDC, IHD or NFH) on γH2AX foci size (*P*<0.001). In FHH, over 40% of γH2AX foci larger than 1 μm^3^ colocalized with telomeres, while only 5.26% and 13.06% of smaller foci demonstrated telomere association in IDC and IHD CMs, respectively (Figure 5G). The observed pattern, considering that telomeres represent less than 0.025% of the genome, is consistent with the preferential localization of larger γH2AX to telomeres. This suggests that DSBs at telomeres, particularly in non-cycling CMs, may be resistant to repair. In contrast, smaller γH2AX foci exhibit a more random distribution across the genome.

### TIFs do not necessarily contain shorter telomeres

Confocal imaging and 3D deconvolution analysis revealed that TIFs contain telomeres of varying lengths, including short, average, and long telomeres, indicating that telomeres within TIFs are not necessarily short in FHH (Figure 6 and Figure S5). Statistical analysis showed a significant difference in TL distributions between telomeres in TIFs and those not associated with γH2AX (out-of-TIF telomeres; Mann–Whitney test, *P*<0.001). The median TL for telomeres in TIFs was significantly higher (Median test, *P*<0.001; Figure 6B), with a greater frequency of telomeres longer than the 95th percentile found in both large and small TIFs (both *P*<0.001, Figure 6C). This suggests that TL alone is not the sole determinant of TIF induction. Additionally, large TIFs (25.00%) were more likely to carry telomere clusters than small TIFs (9.16%) and non-TIFs (0.00%) (Supplemental Table S3). Moreover, GO analysis revealed that telomere maintenance was among the top 45 most overrepresented GO terms for Biological Process, based on the analysis for the co-regulated DEGs in IDC and IHD (Figure 4E). The increase in larger γH2AX foci and their association with TIFs in failing hearts suggest that these foci may function as DNA damage repair centers, accumulate DDR proteins, and potentially facilitate telomere fusion or recombination repair.

### Telomere deprotection and alterations of major DNA-repair factors involved in telomere maintenance in FHH

Human telomeres are protected from DDR by a high-order structure that relies on sufficient telomere and 3’ overhang lengths, as well as shelterin function. This structure is primarily controlled and maintained by TERF2.^7,22^ TERF2 is the main inhibitor of cNHEJ and DNA damage signaling at telomeres by stimulating t-loop formation and directly blocking downstream activators of the ATM DSB signaling cascade. ATR DNA damage signaling at 3’ telomeric overhangs is inhibited by POT1-TPP1 complex, which outcompetes the ssDNA binding protein, replication protein A (RPA) for G-rich overhang binding (Figure 7C).^5,23^ RPA can interfere with telomere capping, as all three RPA subunits feature oligonucleotide/oligosaccharide-binding fold (OB-fold) domains for ssDNA binding. In vitro studies have demonstrated the ability of RPAs to bind and unfold telomeric G-rich 3’ overhangs.^24,25^ Furthermore, it has been proven that oxidative lesions can disrupt the binding of shelterin proteins TERF1 and TERF2 in vitro, impair telomere maintenance, and result in telomere shortening.^26,27^

**Figure 7.**
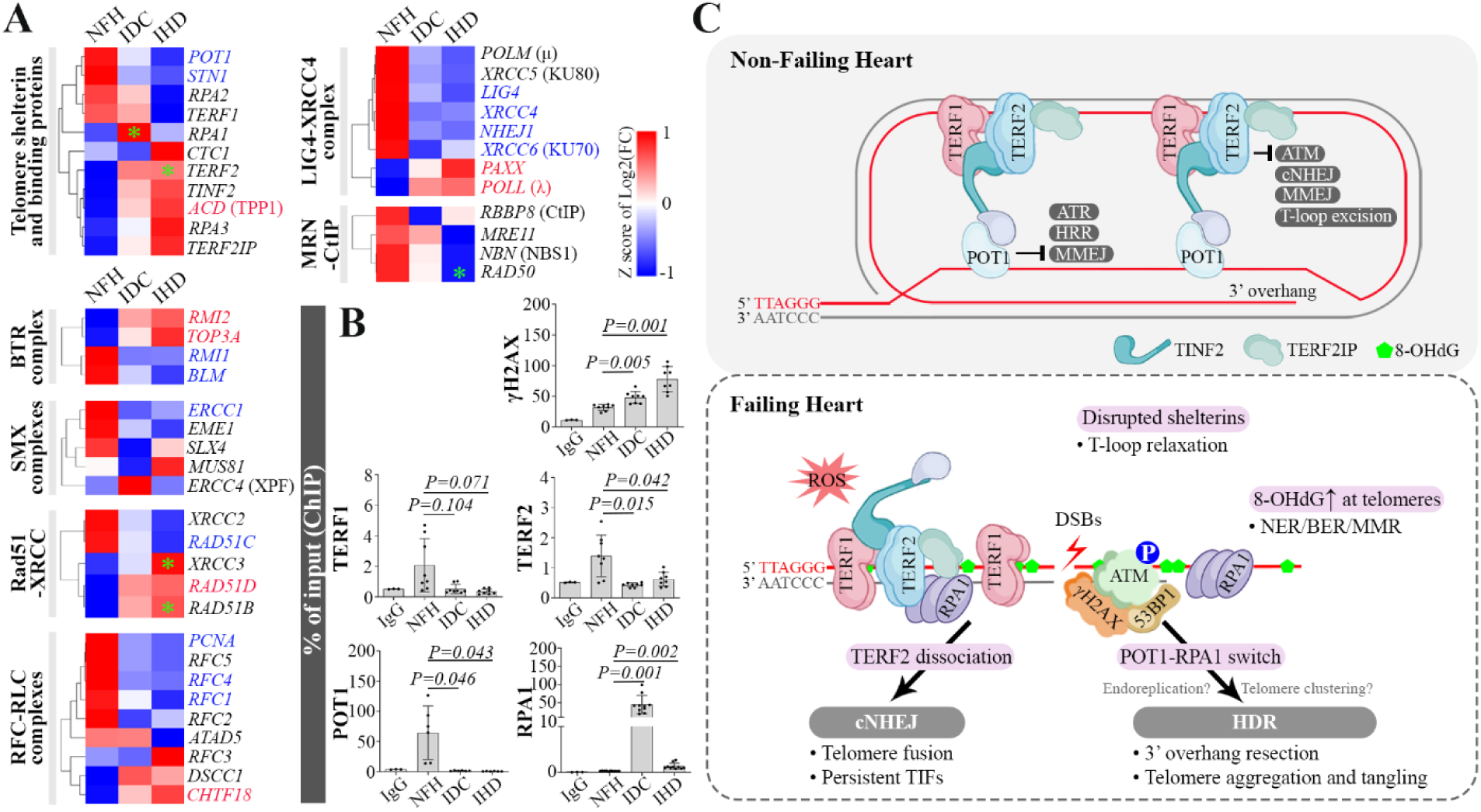
Telomere dysfunction is associated with telomere de-protection. **A**, Heatmaps of telomere maintenance-related genes by high-throughput RNA sequencing from 14 non-failing hearts (NFHs), 14 idiopathic dilated cardiomyopathies (IDCs), and 13 ischemic heart diseases (IHDs) show the log2 fold changes of these targets in IDC and IHD compared to NFH. Differentially expressed targets (DEGs) are highlighted in red if they are co-upregulated and blue if they are co-downregulated in both IDC and IHD; otherwise, they are labeled with green asterisks. In telomere shelterin and binding proteins, the protective single-stranded overhang-binding protein POT1 is decreased in both IDC and IHD, while the DNA damage responsive molecule RPA1 is greatly increased in IDC. **B**, Chromatin immunoprecipitation followed by quantitative polymerase chain reaction (ChIP-qPCR) reveals the binding of TERF2 and POT1 to telomeres is reduced, while the interaction of DNA damage response proteins γH2AX and RPA1 with telomeres is increased. Bar graphs show the quantification of 6-10 hearts for ChIP assays in the NFH, IDC, and IHD groups, and 3 hearts for the IgG control. The results are presented as the percentage enrichment relative to inputs. **C**, The illustration depicts potential pathways and key molecules implicated in the response to telomere-internal DNA damages and the dynamics of the telomere shelterin complex in failing human heart (FHH) cardiomyocytes, elucidating their correlation with telomere alterations in FHH.

Re-analysis of the RNA-seq dataset from human hearts revealed notable changes in telomere-binding proteins in FHH. Specifically, the expression of single-stranded telomere-binding protein POT1 was reduced in FHH, while RPA1 expression increased in IDC. ChIP analysis confirmed a dramatic decrease in POT1 association and an increase in RPA1 association with telomeres, accompanied by elevated γH2AX accumulation in IDC and IHD compared to NFH (Figure 7B). Although TERF1 mRNA levels remained unchanged in FHH and TERF2 was slightly increased in IHD, TERF2 association with telomeres was reduced in both IDC and IHD. Furthermore, IDC and IHD did not show consistent changes in the telomere shelterin interactors that have been proven to directly bind to shelterin proteins (Supplemental Figure S6A and B). These findings suggest that failing hearts undergo telomere deprotection, potentially driven by increased oxidative damage at telomeres, which has been shown to impede the interaction of double and single-stranded binding proteins with telomeres. Moreover, as depicted in Figure 7A, a significant overlap in DDR factor alterations was observed between IDC and IHD, particularly in the LIG4-XRCC4, BLM-TOP3A-RMI1-RMI2 (BTR), SLX1-SLX4-MUS81-EME1-ERCC1-ERCC4 (SMX), and RCF-RLC complexes, which are critical in both cNHEJ and HDR. The transcriptional changes for other potential telomere regulators in FHH are also shown in Supplemental Figure S6B.

## DISCUSSION

The human telomere structure consists of repeating DNA sequences, estimated to range from 5 to 15 kb, followed by a G-rich single-stranded 3’ overhang, which varies between 75 to 300 bases.^1,2,6,28^ The lengths of telomeric duplex and G-tails play a pivotal role in maintaining normal telomere structure and function. Both telomere shortening and 3’ overhang attrition are implicated in cell senescence, which can result from replicative, oxidative, or oncogenic stress. However, the interaction between total TL and 3’ overhang length is intricate, and their respective contributions to senescence and cell dysfunction remain a subject of controversy indicating that both telomeric duplex and 3’ overhang lengths are not fixed, but instead are subject to regulation.^9^

### Alteration of telomere length and 3’ overhang in FHH

While dividing and non-dividing cells can enter senescence in response to stress and experience TL changes during aging, most of our knowledge about cellular senescence and telomere shortening stems from dividing cells. Telomeric DNA shortening, naturally linked to cell division, is generally not considered as a primary driver of aging, senescence, and cell dysfunction in long-lived post-mitotic cells. However, this view is challenged by several reports that have revealed telomere shortening in human and mouse cardiac tissues, even though conflicting data exist regarding the role of telomere shortening in cardiac dysfunction.^10,19,29–32^

Our results suggest that very short telomeres increase in FHH with IDC and IHD, as revealed by Q-FISH analysis of individual CMs. While IDC showed a statistically significant reduction in median TL compared to NFH, this difference was not observed in IHD. Moreover, the relative average TL in FHH, as determined by two different methods, TRF and qPCR analysis of genomic DNA, did not exhibit significant changes. There are several possible factors that may limit the sensitivity and specificity of these methods. First, the DNA samples used for TRF and qPCR analyses were extracted from whole heart tissues. Given that CMs constitute only 25-35% of all cells in the heart^33,34^ and cardiac cellular composition undergoes significant alterations during heart failure progression^35^ due to cardiac hypertrophy, fibrosis, and inflammation, this cellular heterogeneity could greatly influence TL measurements when using DNA from whole heart tissues with heart failure. Secondly, and more importantly, detecting small telomere losses of tens to few hundreds of base pairs may be challenging for these methods when measuring TL under denaturing conditions, considering the proportional relationship between telomere shortening and total TL in human cells. This could partially explain why we discovered that the 3’ G-rich overhang is significantly truncated in FHH compared to NFH. Here, we show that only 3’ G-rich overhang length, and not the TL, exhibits a strong negative correlation with the γH2AX level, which is a DNA damage indicator, in human heart tissues, suggesting that cardiac cells with shorter 3’ G-rich overhangs have more DNA damage accumulation. This is the first time, to our knowledge, that 3’ overhang alteration has been investigated and reported in FHH with IDC and IHD.

### Detection of γH2AX foci and TIFs in CMs of FHH

Our analysis revealed two categories of γH2AX foci within the nuclei of CMs in human hearts. Utilizing image deconvolution and 3D reconstruction with Huygens professional software, we conducted a detailed examination of the intricate structures of γH2AX foci and their spatial relationships to telomeres at the nanometer scale (see Supplemental Methods section). One category manifests as small to medium-sized dot structures, commonly observed in CMs immediately after oxidative challenge, exhibiting a random distribution throughout the nucleus.^36^ In contrast, the other category tends to aggregate, forming larger spatially clustered foci, which notably co-localize preferentially with telomeres to form TIFs, and are predominantly present in CMs from FHH. The clustered foci morphology aligns with previous in-depth spatial analyses of γH2AX foci induced by DSBs, indicating that γH2AX foci comprise spatially distinct nano-domains clustered around DSB sites with median diameters ranging from 150 to 160 nm, median lengths from 375 to 500 nm, and median DNA lengths of 75 kb, as demonstrated through super-resolution microscopy.^37^ These larger γH2AX foci have also been observed in senescent cells and reported as DNA segments with chromatin alterations reinforcing senescence (DNA-SCARS).^38^ They contain DNA repair factors, persist long after DSB induction, and likely harbor unrepairable DSBs.^39^ We confirmed that these larger, clustered foci in FHH contain DNA repair proteins 53BP1 and pATM, but not pATR, in agreement with that the ATM pathway is thought to respond primarily to DSBs, whereas ATR activation requires the formation of ssDNA.

While establishing direct links between 3’ G-rich overhang truncation and TIF formation in individual CMs poses experimental challenges, we found that these TIFs did not necessarily harbor short telomeres. Instead, they often contained longer telomeres, surpassing the 95th percentile, with the higher average TL than out-of-TIF telomere. This suggests that telomere dysfunction can be independent of total TL, consistent with findings reported in both post-mitotic CMs^40^ and proliferation-competent cells.^41^ Our findings, however, do not eliminate the possibility that the longer telomeres in TIFs may result from end-to-end fusions of shortened telomeres. This phenomenon can only be demonstrated and visualized in cultured cells arrested in mitosis through mitotic chromosome preparation,^42,43^ and is difficult to detect in intact tissues, particularly in heart tissues with predominantly post-mitotic CMs.

### Oxidative DNA damage and shelterin protein dynamics in telomere dysfunction

Telomeric regions are highly vulnerable to oxidative DNA damage due to their high guanine content within telomeric repeats.^17,18^ It has been proposed that oxidative modifications to telomeric bases may play a more significant role in telomere loss and telomere-driven senescence than the end-replication problem.^44^ ROS-induced DNA damage contributes to telomere dysfunction in low-proliferative tissues, including the heart.^40,45–48^ Our findings align with this hypothesis, as evidenced by the significant accumulation of 8-OHdG and γH2AX at telomeres in FHH. Key molecules involved in DNA damage repair pathways, including DSB repair and BER, which is primarily involved in resolving oxidized bases, were markedly altered in FHH. Additionally, we observed a significant dissociation of the double-stranded telomere-binding protein TERF2 from telomeres in FHH. Furthermore, our investigation uncovered the dissociation of single-stranded telomere-binding proteins, including POT1, coupled with an enrichment of RPA1 at telomeric regions in FHH, suggesting telomere de-protection and/or t-loop unfolding. While our data showed a significant increase in γH2AX at telomeric repeats in FHH, it is experimentally challenging to determine the initial cause of telomere dysfunction in this context.

### Potential telomere damage repair pathways in FHH

Experimental knockdown or disruption of shelterin components can trigger DDR at telomeres.^2–5^ Conversely, telomere DNA damage often impairs the interaction of shelterin proteins with telomeres.^26,27^ Recent discoveries have indicated that DDR factors play crucial roles in telomere processing, replication, and the establishment of telomere protection.^49–51^

The LIG4 and XRCC4 collaborate to form a ligation complex crucial for repairing DSBs via the cNHEJ pathway. This XRCC4-LIG4 complex is recruited to DNA damage sites through interaction with the XRCC5 (KU80)-XRCC6 (KU70) heterodimer, known as the Ku complex, which serves as a sensor for DSB detection. At the transcriptome level, LIG4, XRCC4, XRCC6, and another essential component for the ligation step of cNHEJ, NHEJ1 (also known as XLF), were significantly reduced in FHH. Conversely, the recently identified paralog of XRCC4 and NHEJ1, known as PAXX, which collaborates with DNA polymerase lambda (POLL) to promote joining of non-cohesive DNA ends, showed an increase in FHH.

Telomeres consist of many kilobases of identical repeats at every chromosome end, making them excellent substrates for HDR. Homologous pairing and strand exchange lead to the formation of DNA intermediates, in which sister chromatids or homologous chromosomes are covalently linked by four-way Holliday junctions (HJs).^52^ In humans, HJs are primarily dissolved by the human Bloom syndrome helicase (BLM)-Topoisomerase IIIα (TOP3A)-RMI1-RMI2 (BTR) complex.^53–55^ The SMX complex, composed of the SLX4 scaffolding protein and the structure-specific endonucleases SLX1, MUS81-EME1, and ERCC1-ERCC4 (XPF), serves a crucial role in resolving recombination intermediates.^56–58^ The activity of BTR must be finely counterbalanced by the SMX complex to prevent DNA breakage and entanglements.^59–62^ In FHH, the mRNA levels of BLM and RMI1 were significantly decreased, while TOP3A and RMI2 were increased. However, the key components of the SMX complex exhibited no changes, except for ERCC1, which encodes the smaller non-catalytic subunit of ERCC1-ERCC4 nuclease. ERCC1 is crucial for NER but not for other DNA repair pathways, including DSB repair.^63^

Extensive and consistent transcriptional alterations were observed in IDC and IHD for the complexes associated with both cNHEJ and HDR, including LIG4-XRCC4, BTR, SMX, and RFC-RLC complexes. This suggests that both pathways are actively involved in the development and progression of FHH. Consequently, DSBs in telomeres can lead to telomere fusion, primarily mediated by cNHEJ, as well as telomere tangling due to unresolved homologous recombination intermediates and telomere end resection mediated by HDR.

### Insights and conclusions

Telomere dysfunction and persistent DDR have been documented in cellular senescence, organism aging, and various human diseases. Our study indicates that these changes also exist in FHH and may play critical roles in the development and progression of human heart failure. Our results suggest, for the first time, that telomere 3’ overhang attrition, rather than overall TL change, is a common characteristic in FHH with IHD and IDC. This alteration is potentially driven by the buildup of oxidative DNA damage in telomeres, leading to the loss of protection by telomere capping proteins, the accumulation of DNA damage, and prolonged DDR activation. Consequently, persistent DNA damage foci, which co-localize with telomeres, increase in CMs of FHH independently of total TL.

It is important to note that DNA repair in adult CMs is prone to errors, as they lack or are deficient in error-free HDR, which requires cell cycle re-entry and the presence of homologous DNA template for precise repair. However, this ability to undergo a full cell cycle is lost in mammalian CMs soon after birth.^64–66^ Growing evidence indicates that these cells may re-enter the cell cycle by undergoing endoreplication and polyploidization in response to stress or during aging.^34,67–69^ This aligns with our findings that FHH has a higher number of polypoid CMs, as well as previous studies showing an increased rate of binucleated CMs in FHH compared to NFH controls.^70^ Heart failure is associated with dysfunctional telomere status rather than absolute telomere loss, which has been detected in human fibroblasts^11^ and age-related myocardial dysfunction.^40^ The intricate interplay among the 3’ G-rich overhang, total TL, and telomere status requires deeper investigation due to its complexity. Further research aimed at uncovering the underlying pathways involved in telomere oxidative damage repair and telomere DDR may provide new avenues and promising therapeutic options for preventing and treating organ dysfunction in various chronic debilitating diseases such as IHD and IDC.

## ABBREVIATIONS

BLM: Human Bloom syndrome helicase
CMs: Cardiomyocytes
cNHEJ: Classical nonhomologous end joining
DDR: DNA double-strand break repair
DSBs: Double-strand breaks
EME1: Essential meiotic structure-specific endonuclease 1
ERCC1: ERCC excision repair 1
ERCC1: ERCC excision repair 1
ERCC4: ERCC excision repair 4
HDR: Homology-directed repair
HRR: Homologous recombination repair
LIG4: DNA ligase IV
MMEJ: Microhomology-mediated end joining
MMR: Mismatch repair
MUS81: MUS81 structure-specific endonuclease subunit
NER: Nucleotide excision repair
NHEJ1: Non-homologous end joining factor 1
RFC: Replication factor C
RLC: RFC-like complex
RMI1: RecQ mediated genome instability 1
RMI2: RecQ mediated genome instability 2
SLX4: SLX4 structure-specific endonuclease subunit
SSA: Single-strand annealing
TL: Telomere length
TOP3A: Topoisomerase IIIα
XRCC4: X-ray repair cross-complementing 4
XRCC5: X-ray repair cross-complementing 5
XRCC6: X-ray repair cross-complementing 6

## Acknowledgments

This research was in part supported by National Institutes of Health grants R01 HL153403 (to FL), RF1 AG072510, R01 HL072166, and R01 HL153614 (to XW).

## Author contributions

FL conceptualized the study. FL and BY designed experiments; BY, LS, XY, and YX performed experiments; BY, LS, and XY analyzed data; FL and BY wrote the initial draft; HX, XW and KBM edited the manuscript. FL and BY supervised the study and approved the final manuscript; FL and XW acquired funding.

